# Effect of head volume conduction on directed connectivity estimated between reconstructed EEG sources

**DOI:** 10.1101/251223

**Authors:** Alessandra Anzolin, Paolo Presti, Frederik Van de Steen, Laura Astolfi, Stefan Haufe, Daniele Marinazzo

## Abstract

Electrical activity recorded on the scalp using electroencephalography (EEG) results from the mixing of signals originating from different regions of the brain as well as from artefactual sources. In order to investigate the role of distinct brain areas in a given experiment, the signal recorded on the sensors is typically projected back into the brain (source reconstruction) using algorithms that address the so-called EEG “inverse problem”. Once that the activity of sources located inside of the brain has been reconstructed, it is often desirable to study the statistical dependencies among them, in particular to quantify directional dynamical interactions between brain areas. Unfortunately, even when performing source reconstruction, the superposition of signals that is due to the propagation of activity from sources to sensors cannot be completely undone, resulting in potentially biased estimates of directional functional connectivity. Here we perform a set of simulations involving interacting sources, and quantify source connectivity estimation performance as a function of the location of the sources, their distance to each other, the noise level, the source reconstruction algorithm, and the connectivity estimator. The generated source activity was projected onto the scalp and projected back to the cortical level using two source reconstruction algorithms, Linearly Constrained Minimum Variance (LCMV) beamforming and ‘Exact’ Low-resolution Tomography (eLORETA). In source space, directed connectivity was estimated using Multi-Variate Granger Causality (MVGC), Time-Reversed Granger Causality (TRGC) and Partial Directed Coherence (PDC), and the estimated connectivity was compared with the imposed ground truth. Our results demonstrate that all considered factors significantly affect the connectivity estimation performance.

## 1. Introduction

Understanding how the joint dynamics of separate brain regions gives rise to function is a fascinating and challenging issue. Several techniques are constantly being developed to investigate these dynamics. EEG signals, due to their high temporal resolution and non-invasiveness, are often employed to investigate how brain activity is modulated in different tasks or conditions [1]–[5]. One of the main issues associated with the EEG signals is the low spatial resolution due to the head volume conduction [6], [7]. It is well known that the electrical activity measured at sensors level is a mixture of the source activity coming from all the sources in the brain (in addition to contributions coming from outside of it). In other words, the spherical geometry of the head and the presence of several tissues with different electrical properties between the cortex and the scalp distort the electric field generated by active neurons so that it is not possible to associate a single brain area to each electrode. The high correlation between signals recorded from neighbouring electrodes at scalp level leads connectivity algorithms to estimate inaccurate patterns including spurious links and to taint results with poor interpretability. Making inferences on connectivity from the EEG signal is still not straightforward [8], [9]. In order to overcome or attenuate the volume conduction problem, several strategies and algorithms have been proposed to estimate source activities from multi-channel EEG recordings [10]. For example, simple spatial filters as the Laplacian can reduce the correlations among scalp-recorded channels induced by the source mixing [6]. Another possibility is given by the Blind Source Separation (BSS) techniques that allow to separate the data into underlying components representing the activity potentially extended networks at the source level. Two algorithms specifically developed for Granger-causal interactions assume that these components follow a multivariate autoregressive (MVAR) model with independent innovation noise [11], [12]. While such approaches allow one to reduce the volume conduction effect, the problem of the interpretability of the results is not completely addressed since directed dynamical influences are estimated between components and not on the cortical brain activity.

Another important choice concerns the connectivity estimator. There are different kind of algorithms and some of them were developed specifically to be less sensitive to artifacts of head volume conduction. Promising results have been obtained using the Phase-Slope Index (PSI) [13] and the Imaginary Coherence (ICoh) [14] but these methods, despite being less sensitive to volume conduction, do not solve the inverse problem and don’t allow precise localisation. Furthermore, their bivariate nature leads in some conditions, to spurious links due to hidden sources. In fact, it is well known how pairwise approaches can lead to false positive detections of connections due to their inability to distinguish a direct interaction between two signals from the influence of a common driver acting on both signals. [15]. Among the directed connectivity estimators, worth of note is the class of multivariate estimators based on the concept of the Wiener-Granger Causality (GC) [16]. These data-driven approaches are computationally simple and require no a priori assumption on the presence or absence of interactions between specific pairs of variables. For this reason we decided to focus on three of them: the classical time-domain measure Multi-Variate Granger Causality (MVGC) [17], its adaptation called Time-Reversed Granger Causality (TRGC) that uses time-reversed data as surrogates for statistical testing [9], [18], and the frequency-domain measure Partial Directed Coherence (PDC) [19].

To achieve interpretable results, the reconstruction of brain sources prior to conducting connectivity estimation is required. To solve the ill-posed (as the number of sources higher than the number of sensors) EEG inverse problem, several algorithms are available. Two of the most commonly used algorithms are the ‘exact’ Low Resolution Tomography (eLORETA) [20], and the Linearly Constrained Minimum Variance (LCMV) Beamformer [21]. Previous studies on real EEG data have already highlighted differences associated with these inverse solutions [22] but additional simulations studies are necessary in order to provide reliable and more specific findings. In other words, different algorithms for the inverse problem solution and for the connectivity estimation could be more or less sensitive to the volume conduction problem, but the evaluation of their performances on real datasets is not possible since an objective ground truth is typically not available. Inverse approaches for extracting cortical waveforms and Granger-based estimators for connectivity measures can be combined to extract and investigate the human brain circuits but a complete evaluation of the volume conduction effect, in terms of demixing quality, in different experimental conditions is still necessary. It is also important to consider the presence of a localization error associated to the forward model used to describe the relationship between activations in the brain and scalp potentials. A suitable forward model for such validations is the ‘New York Head’, which is a highly accurate finite element model (FEM) of the electrical current flow of the average adult human head that is based on the segmentation of a highly detailed magnetic resonance image (MRI) into six different tissue types [23]. The goal of the present study is to identify data analysis pipelines combining source localization approaches and methods for brain connectivity estimation that are able to provide accurate and reliable estimates insensitive to the spurious effects induced by the volume conduction, and thereby allow one to interpret the obtained results in neurophysiological terms. In particular, the present study:

- Demonstrates the possibility to significantly reduce the effect of the volume conduction on the connectivity estimates employing appropriate algorithms as the TRGC;
- Provides guidelines for the employment of the best methods with different spatial distributions of the sources (different depth and relative position);
- Evaluates, which source reconstruction approach, among eLORETA and LCMV, leads to a better performance in this context.

In order to reach these aims, a simulation study was performed starting from the generation of simulated data, which mimic brain source signals with an imposed connectivity pattern. The influence of volume conduction on connectivity estimates was investigated by assigning simulated source signals to different anatomical location in the brain. The results of these simulations allow us to identify the best-performing combination of algorithms for the estimation of the brain activity and connectivity in several realistic conditions.

## 2. Methods

Over the past few decades, different techniques of source localization applied to EEG data were developed to provide a non-invasive estimate of brain activity [24]. Such techniques employ voltage measurements at various locations on the scalp to estimate the current sources inside the brain which best fit these data. Source localization techniques are based on the follow generative model of EEG data:

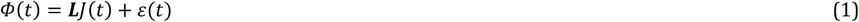

where *ϕ*(*t*) ∈ *R*^*M*^ is the EEG signal measured from *M* scalp locations at time *t*, *J*(*t*) ∈ *R*^3*N*^ is the activity of *N* sources with a 3D orientation in the space, *L* ∈ *R*^*Mx3N*^ is the leadfield matrix summarizing the propagation of the N electrical sources *j* to the EEG sensors and ε(*t*) ∈ *R*^*M*^ is the noise associated to the measures. The lead-field matrix ***L*** contains information about the geometry and the conductivity of all the tissues in the head (between the sensors and the sources) and its computation is well-known as *forward modeling*. The estimation of the sources *j*(*t*) from the measures *Φ*(*t*) contributes to the source reconstruction purpose and it is well-known as *inverse modeling*. The two modeling approaches will be described in detail below.

### 2.1 Forward Problem

The forward problem is solved starting from the electrical activity at source level and calculating the potentials at the sensors (electrodes) level. The result is the scalp activity as a function of the current density (produced by neuronal generators) and describes how the electrical field spreads through the different layers of the head. It depends on the geometry and on the electrical properties of the tissues. The New York Head is an accurate finite element electrical model of the average adult human head [23]. It is based on a highly detailed nonlinear average of T1-weighted structural MR image of 152 adults provided by the International Consortium for Brain Mapping (ICBM) [25]. A detailed segmentation of this average image into six tissue types (scalp, skull, CSF, grey matter, white matter, air cavities) was performed at the native MRI resolution of 0.5 mm3. The suitability of this volume conductor model to serve as an approximation for individual heads was tested by comparison with additional BEMs and FEMs constructed for four subjects.

### 2.2 Inverse Problem

The inverse problem concerns the reconstruction of the brain sources that underlie the measured potentials in electrode space. Because of the difference between the number of sensors and the much higher number of active dipoles in the cortex, the inverse problem solution is not unique. Furthermore, it is very sensitive to small changes in the noisy data and also depends on the choice of the reference electrode. The accuracy of the source reconstruction is affected by a high number of factors including the head model errors, the source-modelling errors and EEG noise (instrumental or biological) [26]. Several algorithms were developed to solve the inverse problem. In the present study, we focused on two methods: i) the Linearly Constrained Minimum Variance Beamformer (LCMV) and ii) the ‘Exact’ Low Resolution Tomography (eLORETA).

#### Linearly Constrained Minimum Variance (LCMV)

Linearly constrained minimum variance filtering (LCMV) [26] is a spatial filtering method that lets brain activity coming from a specific location pass, while attenuating activity originating at other locations. The output of the filter is an estimate of the power of the electrical field generated by the neurons within a restricted area of the brain. The spatial pass-band of the filter depends on the dimension of that area, thus the higher the desired resolution the smaller required pass-band. A map of neural power as a function of location is obtained by designing multiple spatial filters, each with a different pass-band, and depicting output power as a function of pass-band location. This spatial filtering approach falls within the general category of beamforming. It is known that the signal at each location in the brain consists of the three dipole moments, so that three spatial filters for each location are required. The N x 3 matrix ***W***(q_0_) represents the transfer function of the filter for the narrowband volume element Q_0_ centered in q_0_. The output of the filter, *J*, is the inner product between W(q_0_) and *ϖ*.

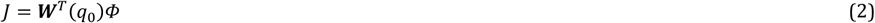

Under ideal conditions, the transfer function of the filter has to satisfy two conditions:

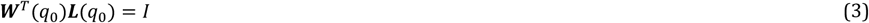

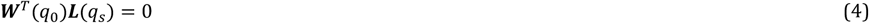

As this cannot be achieved under general conditions, *eq*. 4 is replaced by the condition that the variance of the filter output (*eq*. 5) is minimal.

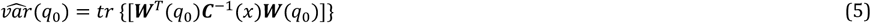

The optimal filter is given by:

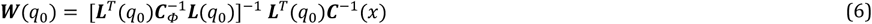

The variance of the filter output can then be simplified as

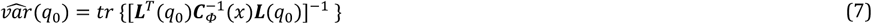

where the sensor-space covariance is:

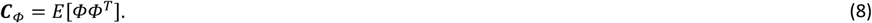

The optimal filter *W*(*q*_*s*_) has a large output in *q*_*s*_ only if there is a significant energy originating from there. To localize the electrical activity of the brain sources, the variance of the LCMV filter output is evaluated as a function of location within the volume of the brain, normalized by the LCMV filter output on a reference (noise) data segment. Regions of large relative variance are presumably active, while regions with small relative variance can be considered inactive. Nevertheless, in the present study, the goodness in terms of localization is not it is not of main interest. We only considered the estimated source time series (filter output) to assess connectivity patterns between them. Factors that may influence the accuracy of the LCMV are:

- The pass-band of the filter, indicating the spatial resolution. The spatial extent of the pass-band depends on the transfer matrices ***L**(q)*, which in turn depend on the number of electrodes, their distribution, and source location.
- The SNR, because of the variance minimization procedure used to determine the spatial filters. In this context SNR has to be thought of not as ratio of the signal power to the noise power, but rather as the variance of the source divided by the variance of the noise.

#### ‘Exact’ Low Resolution Tomography (eLORETA)

‘Exact’ Low Resolution Electromagnetic Tomography (eLORETA) [27] is a linear inverse method characterized by spatially smooth current density. In the most general case, linear solutions to the EEG inverse problem are of the following form:

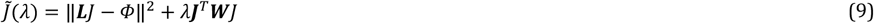

where λ represents the Tikhonov regularization parameter which can be estimated through the general cross validation approach [28], and where ***W*** is a symmetric positive definite weight matrix. The idea of eLORETA is to find an appropriate ***W*** matrix in *eq*. 9 such that the solution has zero localization error for all single point sources in the brain [27]. These weights are obtained from the following expression:

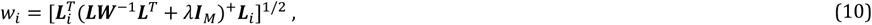

where *w*_*i*_ for *i* = 1,…, *N* (number of voxels) are the diagonal elements of the weight matrix ***W***, *L*_*i*_ ∈ *R*^*Mx1*^ represents the *i-th* column of lead field matrix ***L*** and the symbol + refers to Moore-Penrose pseudoinverse. The solution to (*eq*. 10) can be found by iterating four steps. First, we have to initialize the diagonal matrix ***W*** with *w*_*i*_ = 1, for *i* = 1,…, *N* and then compute:

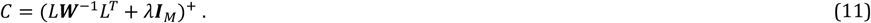

Holding C fixed, we compute new weights for all the dipoles *i* = 1,…, *N*:

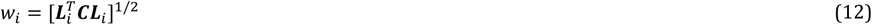

and then we return to *eq*. 11 until convergence. Once the *w*_*i*_ have been estimated, the eLORETA solution is given by the following expression:

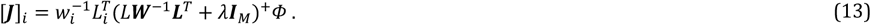

It has been suggested that eLORETA solution achieves exact localization to single test point sources under ideal (no-noise) conditions, outperforming all other linear solutions on both simulated and real EEG data in this respect [29]. However, in the presence of two or more sources (thus, in any setting involving source interaction), this property does not hold anymore.

### 2.3 Multivariate Directed Connectivity Estimation

#### Multivariate Granger Causality (MVGC)

The concept of Granger causality [16], [30] is based on the predictability of time series. Namely, if a time series *X_2_*(*t*) contains information that improves the predictability of future values of another time series *X_1_*(*t*) above and beyond what can be predicted on the basis of *X_1_*(*t*) alone, then *X_2_*(*t*) is said to Granger-cause *X_1_*(*t*). In other words, if the prediction error decreases by adding the past values of *X_2_*(*t*) to a regression model for predicting *X_1_*(*t*), we can assume that *X_2_*(*t*) Granger-causes *X_1_*(*t*). In the BIVAR (bivariate vector-autoregressive) formulation, this notion is described as follows:

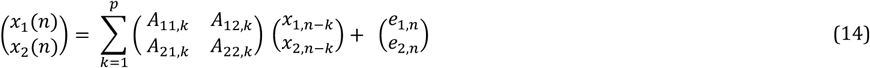

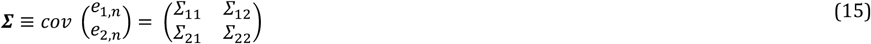

At this point, one can perform a *full regression* (*eq*. 16), using both time series, and a *reduced regression* (*eq*. 17), using only the target time series:

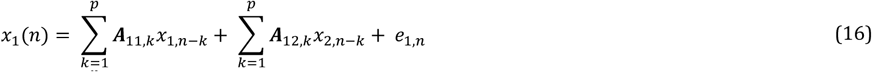

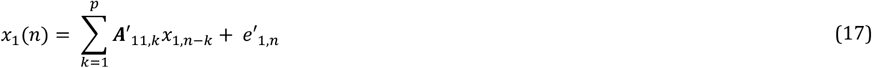

In the full regression, the dependence of *X*_*1*_ on the past of *X*_*2*_, in addition to its own past, is encapsulated in the coefficients ***A***_*12*,*k*_. There is no dependence between *X_1_* and *X_2_* if the coefficients are null for all lags *k*, ***A***_*12,1*_ = ***A***_*12,2*_ =… = ***A***_*12,p*_ = *0*. Prediction error estimation is based on full and reduced regression residuals. In particular ***Σ***’_*11*_ ≡ *var*(*e’*_*1,n*_) is the residual variance in the case of reduced regression and ***Σ***_*11*_ ≡ *var*(*e*_*1,n*_) is the residual variance in the case of full regression. Pairwise time-domain Granger causality is defined as

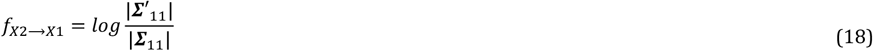

The value of *f*_*x2*→*x1*_ is equal to 0 if there is no GC between the time series and their variance ratio is 1. If a dynamical influence from *X_2_* to *X*_*1*_ exist, the value of *f*_*x2*→*x1*_ is greater than zero. Let us suppose to have joint dependencies between *X*_*1*_ and *X*_*2*_ and a third set of variables, e.g. *X3*, then spurious influences may be reported. Spurious connections can be detected even when there is no direct influence *X*_*2*_→*X*_*1*_ but there are (possibly lagged) dependencies of *X*_*1*_ and *X*_*2*_ on *X*_*3*_. To overcome this problem, Barnett and Seth propose a different way to compute GC, introducing the so called *Pairwise Conditional Granger Causality (PWCGC)*, which conditions out common dependencies between variables before estimating pairwise GC scores, provided such dependencies are present in the data [31]. The MVAR model is again expressed in the form of full regression (*eq*. 19) and in the form of reduced regression (*eq*. 20), as:

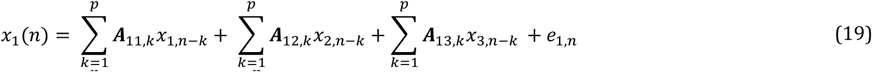

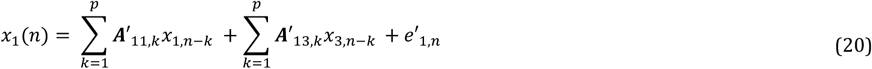

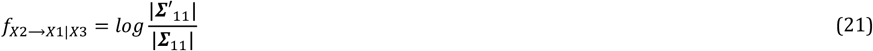

Here, *F*_*X2→X1|X3*_ may be read as “the degree to which the past of *X*_*2*_ helps to predict *X*_*1*_, over and above the degree to which *X*_*1*_ is already predicted by its own past and the past of *X*_*3*_”. In our simulation study, we are going to use this approach. Additionally it is worth noting that we use the state-space formulation of Granger causality, which eliminates the bias due to the fact that the reduced model is VARMA (Vector Auto Regressive Moving Average) and not VAR [32].

#### Time reversed Granger causality

Granger-causal estimators are prone to detect spurious influences not only in the presence of hidden common drivers but also in the presence of additive correlated noise [9], [13], [18], [33], [34]. Correlated noise is a ubiquitous property of EEG data, which are by their very nature linear mixtures of contributions from different sources. Since this mixing process cannot be fully undone using source imaging techniques, it poses a serious problem for EEG-based brain connectivity analysis using GC. To overcome the problem of spurious connectivity for mixed data, Haufe et al. proposed time-reversal [9], [34]. The intuitive idea behind this approach is that, if connectivity is defined based on temporal delays, directed influence should be reduced (if not reversed) if the temporal order is reversed. This is in contrast to the observation that two signals that are correlated but Non-Interacting often appear spuriously connected no matter whether GC is applied on the original or time-reversed data. If, however, GC estimates obtained on original and time-reversed data are contrasted with each other, the instantaneous influence of volume conduction can be removed, and the false detection of connectivity can be avoided. GC is defined based on the *Granger-scores* defined in *eq*. 18, where *Fx_1_*→*x_2_* is the direct influence from *x_1_* to *x_2_*, and it requires that the residual variance of the restricted model should be smaller than the one in the case of full model [13]. When time-reversing the data, we denote the residual covariance matrix of the time-reversed process (full model) by:

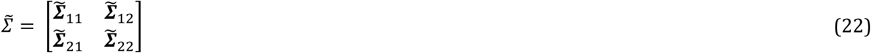

As for the original GC, we define the dynamical influence as

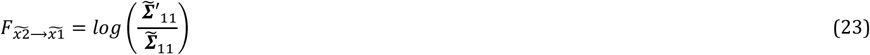

Finally, Time-reversed GC is given by the difference between the net GC scores obtained on the original and time-reversed GC:

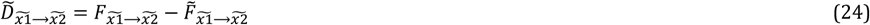

Using the above definitions, the validation of a Granger causal influence that cannot be explained by a mixture of independent sources can be performed according to the following criterion, named Conjunction-based time-reversed GC:

- the directionality of GC is required to flip for time-reversed signals. The connection is regarded as significant if both GC values (with original and reversed data), are significant: 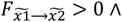 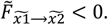. This is the definition adopted in the present paper.

Other criteria, less stringent than this one, are discussed in [18]. Simulations have shown that TRGC leads to a reduced number of false connections, compared to original GC and its variants [9], [18], [33]–[35].

Theoretical work presented in [18] has moreover shown that:

- The application of time reversal to any connectivity measures that is based on second order statistics - which, besides GC and pairwise-conditional GC also includes its direct extension to frequency domain (spectral GC) and the popular frequency-domain measures partial directed coherence (PDC) and directed transfer function (DTF), among others - prevents the spurious detection of connectivity on mixtures of independent sources that would otherwise be highly likely.
- The application of time reversal to Granger causality (that is, the use of TRGC) is guaranteed to always yield the correct direction of interaction for systems that do not contain causal loops, and are noise-free.

#### Partial directed coherence (PDC)

Partial Directed Coherence (PDC) [36] is a spectral measure to assess the dynamical influence between signals within a multivariate dataset. It is basically a frequency version of the concept of Granger causality [37]. PDC is defined as follows:

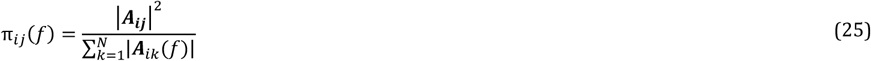

PDC values fall in the range [0, 1], where *π*_*ij*_*(f)=0* stands for the absence of a direct influence from xi to xj at the considered frequency *f*. PDC only estimates the direct influence between two signals, thus discounting for indirect effects of other channels in a similar way as pairwise-conditional GC. The definition has been subsequently refined with the introduction first of a row-wise normalization [38]:

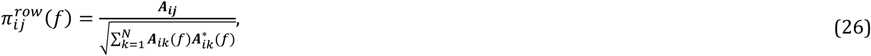

then of a quadratic version, so that the new squared and normalized PDC can be interpreted as the portion of the *ith* signal power density due to the *j*^*th*^ one:

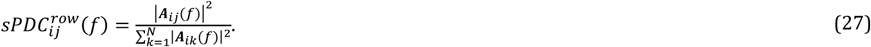

### 2.4 Statistical assessment of significant connections

The standard way to assess the statistical significance of Granger scores is a likelihood ratio test, which can be derived from large-sample theory [39]. If dim(X_1_) = n_x1_, dim(X_2_) = n_x2_ and dim(X_3_) = n_x3_ (with nx1 + nx2 + nx3 = n) then the difference in the number of parameters between the full model and the nested reduced model (see *eq*. 20) is just d ≡ p nx1 nx2. Thus, under the null hypothesis of zero Granger-causal influence, the GC estimator scaled by sample size, (m − p) *F*_*X2*→X1|X3_(u), has an asymptotic _χ_2 distribution. Under the alternative hypothesis, the scaled estimator has an asymptotic noncentral - _χ_2 (d; ν) distribution, with non-centrality parameter ν = (m − p) *F*_*X2*→X1|X3_(u) equal to the scaled actual influence (which may, for the purpose of constructing confidence intervals, be replaced by its estimator). Similarly, it was demonstrated that the squared PDC estimator tends to a Gaussian distribution in the non-null case and to a χ2 distribution in the null case. This assumption led to the development of a new approach, the asymptotic statistic, which allowed the derivation of the probability distribution of the null-case squared PDC estimator (the χ2 distribution), by knowing its asymptotic variance [40], [41]. Note that these standard statistical tests are only capable to distinguish actually present GC/PDC effects from results obtained due to random signal fluctuations in the absence of GC/PDC. They are *not* capable of distinguishing actual GC/PDF effects that are due to genuine time delayed interaction from actual GC/PDC effects that are solely due to additive mixed noise in the absence of genuine time-delayed interaction. To test for the latter, the statistical significance of TRGC (or, time-reversed PDC) needs to be established. For difference-based TRGC, this can be achieved by testing whether (eq. 28) is significantly different from zero using non-parametric approaches like the bootstrap. In this work, we focus on conjunction-based TRGC and we used an alpha of 0.05, FDR corrected.

### 2.5 Simulation Framework

The simulation study developed for investigating the effects of the volume conduction on connectivity estimation accuracy and reliability is composed by the following main steps:

- Generation of brain signals with an imposed connectivity pattern
- Forward problem solution
- Inverse problem solution
- Connectivity estimation
- Performance evaluation

An overview of the simulation framework, with all the considered factors, is shown in *fig*. 1.

**Figure 1.**
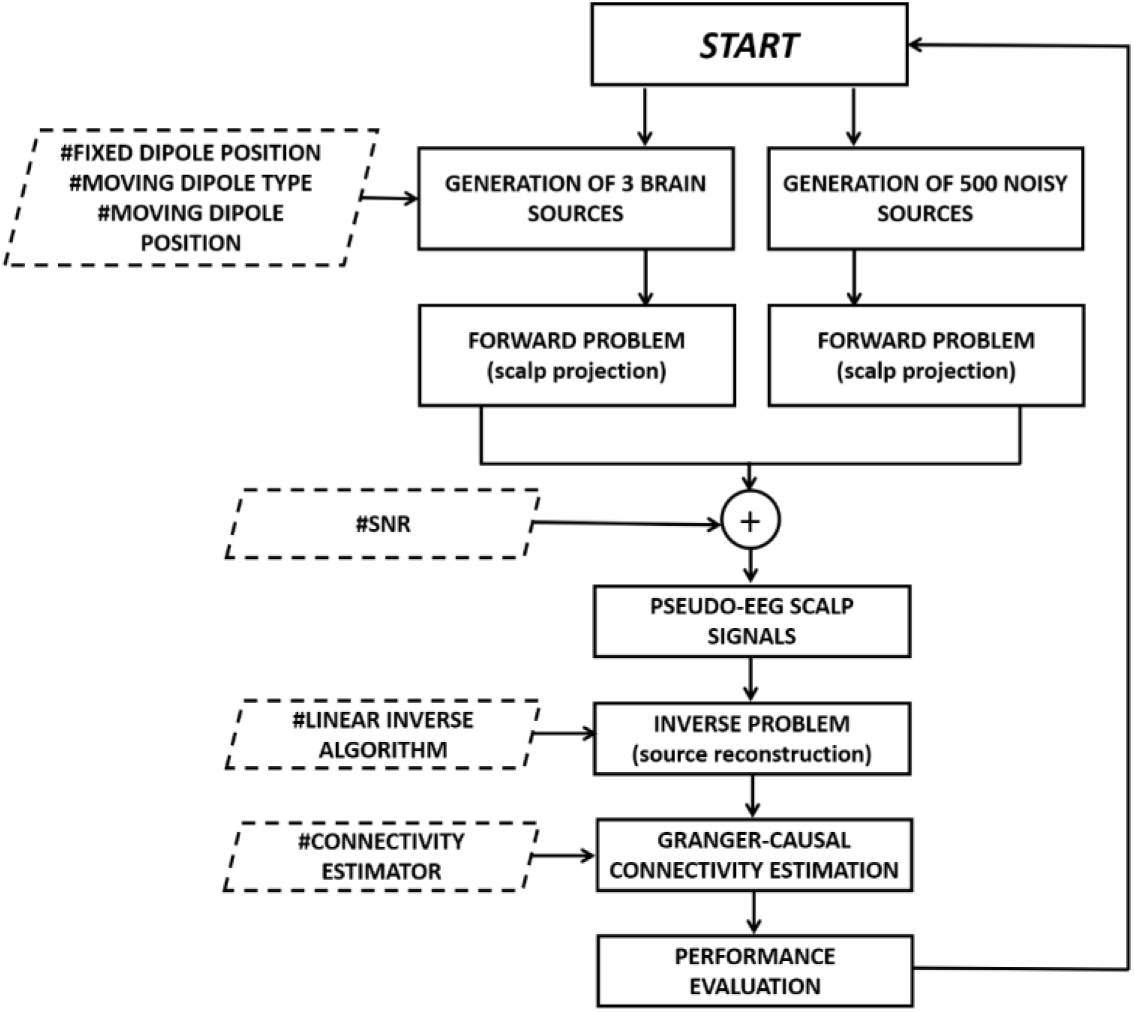
*Block diagram reporting the main steps of the simulation framework*.

### 2.6 Simulated time series generation

Brain signals were generated using a multivariate autoregressive (MVAR) model with order 2 as generator filter. We simulated three time series and only one connection. Both, the autoregressive components and the off-diagonal elements of the coefficients matrix were randomly chosen within the range [0.3 1]. The three different time series will be called *Sender*, *Receiver* and *Non-Interacting dipole*, to indicate, respectively, the driving dipole, the receiving dipole, and the independent dipole. Each of them represents an active source contributing to the simulated EEG scalp potentials. In order to simulate an experimental condition as realistic as possible, we also generated 500 pink noise signals representing the background brain activity.

### 2.7 Simulated time series location

Brain activity was modelled with 1006 electric equivalent dipoles, equally distributed within the brain. Using the New York Head model, we obtained the dipole positions by subsampling the 75000 MNI coordinates available in the ICBM152 model. In the panel a of *fig*. 2 we showed all the 1006 possible dipole locations.

**Figure 2.**
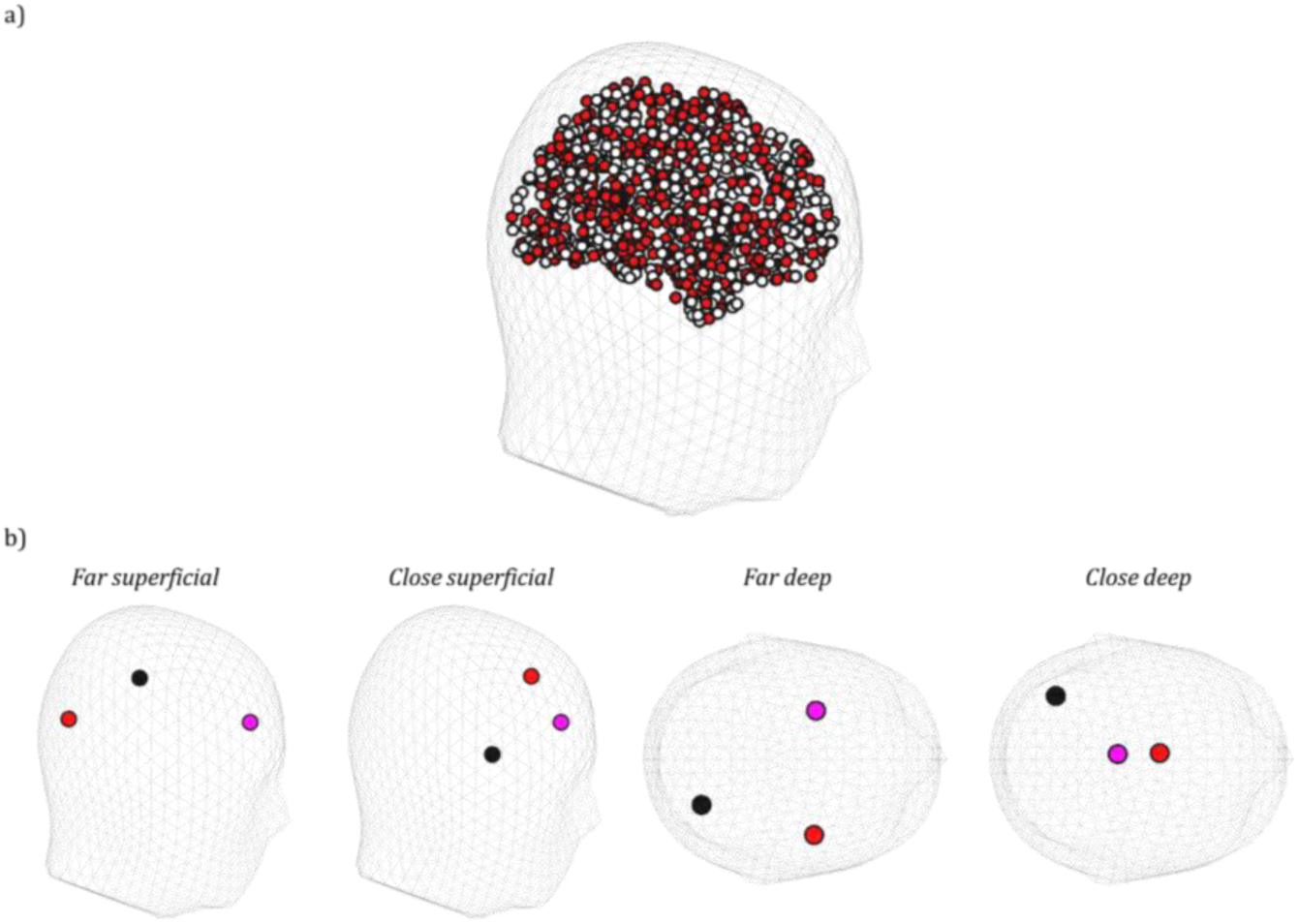
*Panel a) shows the 1006 locations in which activity was modelled. Red circle represents an example of the 500 locations associated with the brain noise activity. Panel b) presents the four conditions for the two fixed active dipoles, which are the red (Sender) and the purple (Receiver) one. The black circle represents the Non-Interacting dipole (noise)*.

For each simulation, we fixed the position of two active dipoles on four possible conditions (represented in *fig*.2b) defined from the combination of the following factors:

- Depth of the dipoles: “superficial” (distance from the origin >6.5 cm) or “deep” (distance from the origin <6cm);
- Distance between the dipoles: “far” (relative distance >8cm) or “close” (relative distance <5cm).

The third dipole (*moving dipole*) instead, change its location on the 1004 possible remaining positions. Two different cases were analysed by changing the moving dipole. In the first case, it is the Non-Interacting dipole, thus the connection is fixed. In the second case, the moving dipole is the Receiver; thus, the relative locations of Sender and Receiver varies across repetitions. The 500 additional noisy elements were randomly distributed within the brain.

### 2.8 Pseudo-EEG signal generation

After the signals generation, the time series representing both the source activity and the noise, were projected onto 108 EEG electrodes defined by the New York Head model, and summed according to following equation:

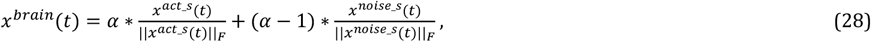

where *x*^*act_s*^ and *x*^*noise_s*^ are the projections of the active sources signals and of the brain noise sources activity respectively, and where ||*x*(*t*)||_F_ is the Frobenius norm of the multivariate time series x(t) (the square-root of the sum of the squared activity across time and space). The parameter *α* thereby defines the signal-to-noise ratio. Given *α*, the corresponding SNR in decibels (*dbs*) is:

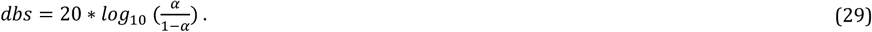

Finally, in order to simulate the measurement noise, spatially and temporally uncorrelated signals are added to *x*^*brain*^(*t*) with an imposed α equal to 0.9. The overall pseudo-EEG data is defined from the following equation:

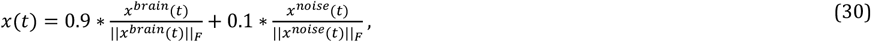

where *x*^*noise*^ is the white uncorrelated noise.

### 2.9 Source reconstruction and directed connectivity estimation

The simulated pseudo-EEG signal was projected onto the cortical surface using two different inverse problem solutions: LCMV and eLORETA. The regularization parameter to be set in the eLORETA algorithm was chosen by means of a cross-validation approach. In cortical source space, directed connectivity according to MVGC, TRGC and PDC was estimated at the locations of the three simulated active dipoles, and the statistical significance of the estimated connections was assessed.

### 2.10 Performance parameters

The quantitative evaluation of the accuracy in signals reconstruction and connectivity estimation was performed by means of three parameters: the False Positive Rate (FPR), the False Negative Rate (FNR) and the Area Under ROC Curve (AUC). Such parameters were computed by comparing the estimated connectivity pattern with the imposed ground-truth. A false positive (FP) is an estimated (statistically significant) connection that is not present in the simulated data, while a true negative (TN) is an absent simulated connection that is correctly estimated as being absent. The FPR (see *eq. 31*) is the number of false positives normalized by the number of absent connections. The FPR is thus defined as in the follows:

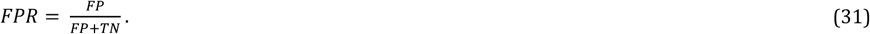

The FNR quantifies the percentage of missed (not statistically significant) connections (referred to as false negatives, FN) that are actually present in the simulated data relative to the total number of actually present simulated connections. The latter number is given as the sum of false negatives and true positives (TP, referring to actually present connections that are also estimated to be present). The FNR is thus defined as follows:

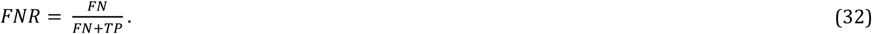

In this study, the total number of possible connections is six (2 possible directions for three distinct pairs of variables). As only one interaction was modelled, FN+TP equals one, while the number of absent connections (FP + TN) is equal to five.

The AUC is a measure of binary classification accuracy, which is applied here to the problem of distinguishing between interacting and Non-Interacting signals. It takes into account both the FPR and FNR across the entire range of all possible thresholds for the connectivity measure; therefore, it is independent of a specific significance level. The AUC is bounded between 0.5 (chance-level class separation) and 1 (perfect class separation) and was derived from the Wilcoxon-Mann-Whitney test [42].

### 2.11 Statistical Analysis

In order to statistically evaluate the accuracy of the employed algorithms in reconstructing the sources activity and estimating brain networks, a four-way ANalysis Of VAriance (ANOVA) was computed. The main within factors were:

- the fixed dipoles position (DIP_POS) with 4 levels: Close Deep, Close Superficial, Far Deep, Far Superficial;
- the adopted inverse methods (L_INV_METH) with 2 levels: eLORETA, LCMV;
- the connectivity estimator (EST_TYPE) with 3 levels: MVGC, PDC, TR_GC;
- the signal-to-noise ratio (SNR) defined by 3 levels of α: 0.5, 0.7, 0.9 (corresponding to SNR equal to 0, 7 and 19 dB respectively) that in the next will be identified as “low”, “medium” and “high” value of SNR.

The dependent variables were the three introduced performance parameters (FPR, FNR and AUC) averaged on the 1004 possible location of the moving dipole. The simulation was repeated 100 times for each experimental condition. Additionally, a post hoc analysis was performed in order to highlight the significant comparisons between the various level of the included factors and their interaction, using Tukey's range test.

### 2.12 Topographical visualization of the results

As described in the previous paragraph, the ANOVA investigates the performance parameters averaged for more than one thousand possible locations of the moving dipole. In order to obtain a detailed overview on the variations of the estimate accuracy as function of the position of the moving dipole, we averaged the parameters on the 100 iterations and reported the obtained results in 3D brain maps. The color of each one of the 1004 dipoles codes for the value of the FPR. We do not report the maps obtained for the false negatives because their amount is always very low (less than 5%).

With the aim to summarize the complex information contained in the brain maps, we also calculated the FPR as function of the distance between Sender and Receiver as well as between Sender and Non-Interacting dipole for each SNR level, inverse approach, and connectivity estimator. The position of the fixed dipole (either Receiver or Non-Interacting dipole) in these analyses was *far* and *superficial*.

## 3. RESULTS

### 3.1 Statistical analysis

The results of the four-way ANOVA computed separately for the three performance parameters are reported in Table I. A four-way ANOVA consists of fifteen separate multiple tests (four main effects, six two-way interactions, four three-way interactions, and one four-way interaction). Therefore, a correction for multiple comparisons (Bonferroni-Holm for example) was performed.

**Table 1.** Results of the four-way ANOVA (F values) computed considering as dependent variables FPR, FNR and AUC and as within main factors the type of inverse algorithm (L_INV METH), the connectivity estimator (EST_TYPE), the SNR and the position of the fixed dipoles (DIP_POS). In the column “Factors”, the degrees of freedom are also reported.

| *Factors* | *FPR* | *FNR* | *AUC* |
| --- | --- | --- | --- |
| L_INV METH (1,99) | **86,25** $p=1*10^{-5}$ | **68,27** $p=1*10^{-5}$ | **41,43** $p=1*10^{-5}$ |
| EST_TYPE (2,198) | **1686,5** $p=1*10^{-5}$ | **8,45** $p=0,0003$ | **1652,6** $p=1*10^{-5}$ |
| SNR (2,198) | **357,14** $p=1*10^{-5}$ | **67,42** $p=1*10^{-5}$ | **570,29** $p=1*10^{-5}$ |
| DIP_POS (3,297) | **930,51** $p=1*10^{-5}$ | **62,13** $p=1*10^{-5}$ | **1292,1** $p=1*10^{-5}$ |
| L_INV METH x EST_TYPE (2,198) | **14,55** $p=1*10^{-5}$ | **15,16** $p=1*10^{-5}$ | **13,03** $p=1*10^{-5}$ |
| L_INV METH x SNR (2,198) | **23,68** $p=1*10^{-5}$ | **101,48** $p=1*10^{-5}$ | **2,5** $p=0,084$ |
| EST_TYPE x SNR (4,396) | **91,06** $p=1*10^{-5}$ | **45,73** $p=1*10^{-5}$ | **74,73** $p=1*10^{-5}$ |
| DIP_POS x L_INV METH (3,297) | **82** $p=1*10^{-5}$ | **86,54** $p=1*10^{-5}$ | **15,14** $p=1*10^{-5}$ |
| EST_TYPE x DIP_POS (6,594) | **225,55** $p=1*10^{-5}$ | **5,69** $p=1*10^{-5}$ | **181,71** $p=1*10^{-5}$ |
| DIP_POS x SNR (6,594) | **84,16** $p=1*10^{-5}$ | **90,47** $p=1*10^{-5}$ | **59,74** $p=1*10^{-5}$ |
| L_INV METH*EST_TYPE*SNR (4,396) | **34,76** $p=1*10^{-5}$ | **27,24** $p=1*10^{-5}$ | **18,96** $p=1*10^{-5}$ |
| L_INV METH*EST_TYPE*DIP_POS (6,594) | **23,43** $p=1*10^{-5}$ | **24,61** $p=1*10^{-5}$ | **29,39** $p=1*10^{-5}$ |
| L_INV METH*SNR*DIP_POS (6,594) | **14,54** $p=1*10^{-5}$ | **79,11** $p=1*10^{-5}$ | **54,39** $p=1*10^{-5}$ |
| EST_TYPE*SNR*DIP_POS (12,1188) | **3,42** $p=1*10^{-5}$ | **28,93** $p=1*10^{-5}$ | **3,98** $p=1*10^{-5}$ |
| L_INV METH x EST_TYPE x SNR x DIP_POS (12,1188) | **47,67** $p=1*10^{-5}$ | **17,3** $p=1*10^{-5}$ | **42,27** $p=1*10^{-5}$ |

All factors and all interactions between factors have a significant effect on the FPR, FNR and AUC. In the following, we show a graphical depiction of the means of the four-way interaction factor (L_INV METH x EST_TYPE x SNR x DIP_POS) for each investigated performance measure.

#### False Positives Rate

Figure 3 shows means obtained for the FPR for different levels of SNR (α) and dipole positions when specific algorithms for the inverse solution and connectivity estimation are employed.

**Figure 3.**
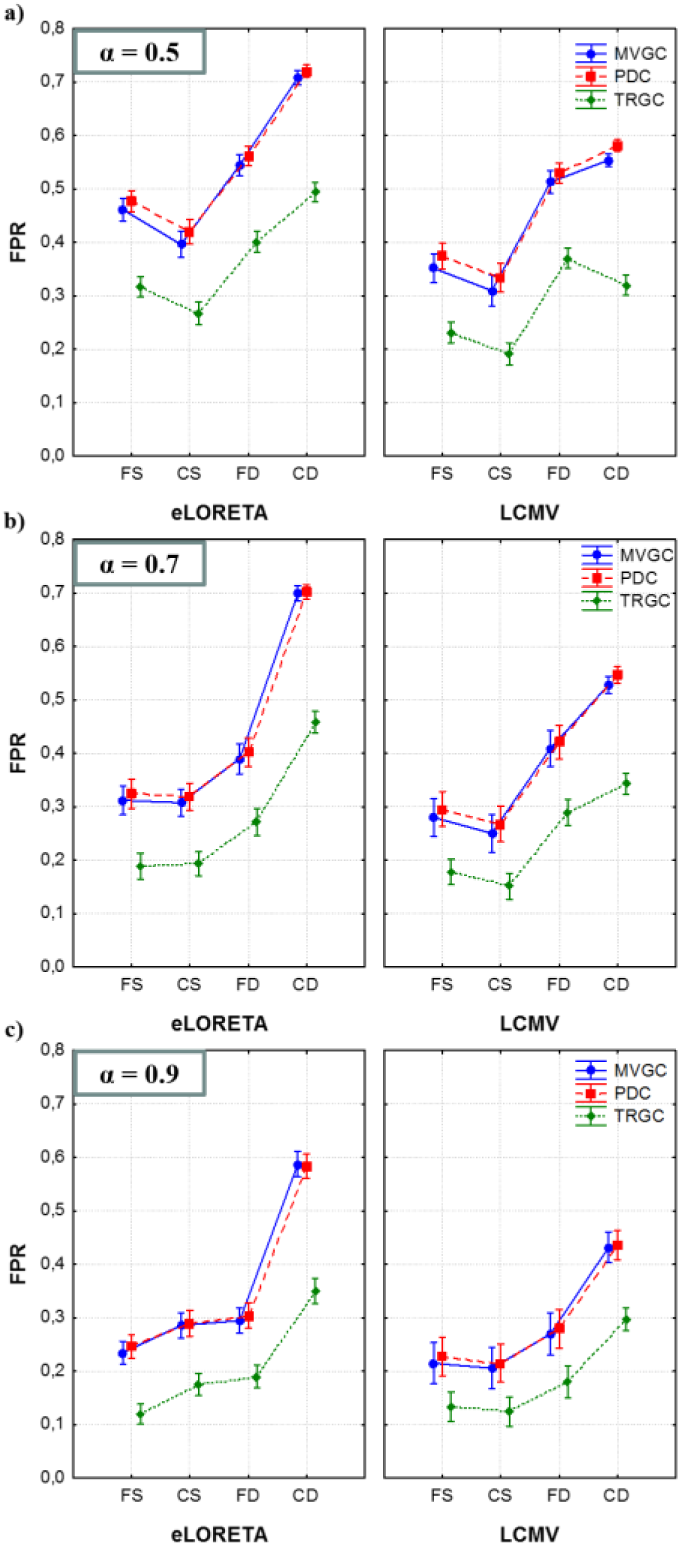
Means associated with the four-way interaction factor (L_INV METH x EST_TYPE x SNR x DIP_POS) of the ANOVA performed on the FPR. Each panel corresponds to a specific value of the SNR parameter α: 0.5 (panel a), 0.7 (panel b), 0.9 (panel c). For each panel, there are two graphs associated with the two different inverse solutions: eLORETA on the left and LCMV on the right. X-axes report the levels of the factor DIP_POS and the colours code for the three connectivity estimators. Whiskers represent 95% confidence intervals. Each panel depicts results obtained for one SNR level.

These graphs show how the two different inverse methods and the location of the fixed dipoles influence the amount of false positive connections when the estimation is performed with the three different connectivity estimation algorithms for different levels of SNR.

##### Connectivity Estimator

As expected, we found that the amount of false positive decreases when the connectivity pattern is extracted by means of TRGC. FPR associated with the TRGC is significantly lower (Tukey test) with respect to the other two methods independently of the dipoles position, the SNR level and to the inverse algorithm (see all the subplots). MVGC and PDC do not show significantly different results for each condition, and the number of estimated spurious connections is not significantly different.

##### Inverse Algorithm

For each panel, we can compare the performance associated with the different inverse solutions comparing the two subplots. Regardless of the SNR, the LCMV algorithm (on the right) for source reconstruction has globally better performance than eLORETA (on the left) for all the three SNR values. The post hoc analysis reveals a significant increase of the FPR for eLORETA, compared to LCMV, in all the considered conditions of SNR, dipoles position, and connectivity estimator. Only in the most advantageous configuration, when α equals 0.9, indicating high SNR, and the linked dipoles are in the *Far/Superficial* configuration, such difference is not significant. In the worst case, corresponding to the *Close/Deep* configuration, the FPR is considerably high, especially for the eLORETA reconstruction, where it exceeds 70%, and the difference between the performances of the two inverse methods appears to be emphasized.

##### Fixed dipoles position

It is worth to note how performance critically depends on the position of the fixed dipoles. Independently of the employed inverse algorithm and connectivity estimator, the ANOVA suggested that when they are located deep in the brain the amount of false positives significantly increases. When the fixed dipoles are superficial and α is equal 0.7, the relative distance (close/far) does not have a significant influence on the FPR. For the higher SNR levels, the ANOVA highlighted a significant increase of the FPR from 10% to 20%, suggesting that the optimal condition for the source reconstruction is given by far and superficial dipoles. For the very low SNR value of 0.5, for all the L_INV METH and EST_TYPE levels, the statistical test revealed a significant decrease of FPR in the *Close/Superficial* case relative to the *Far/Superficial* case.

##### SNR

In all considered conditions, the test indicates a significant improvement of performance when the simulated SNR is higher. More in detail, when the SNR level is 0.9, the amount of false positives is less than 30% in all the cases except for the *Deep/Close* condition. The analysis of FPR suggests that the best combination of factors is given, for all the considered SNR levels, by: i) dipoles located superficial in the brain and not too close; ii) LCMV as algorithm for the inverse problem solution and iii) TRGC as connectivity estimator. Only in this case the percentage of false positives reached low values (around 10% for SNR equal to 0.9).

#### False Negative Rate

The graphs in Figure 4 depict the means of the four-way interaction factor (L_INV METH x EST_TYPE x SNR x DIP_POS) obtained for the FNR index.

**Figure 4.**
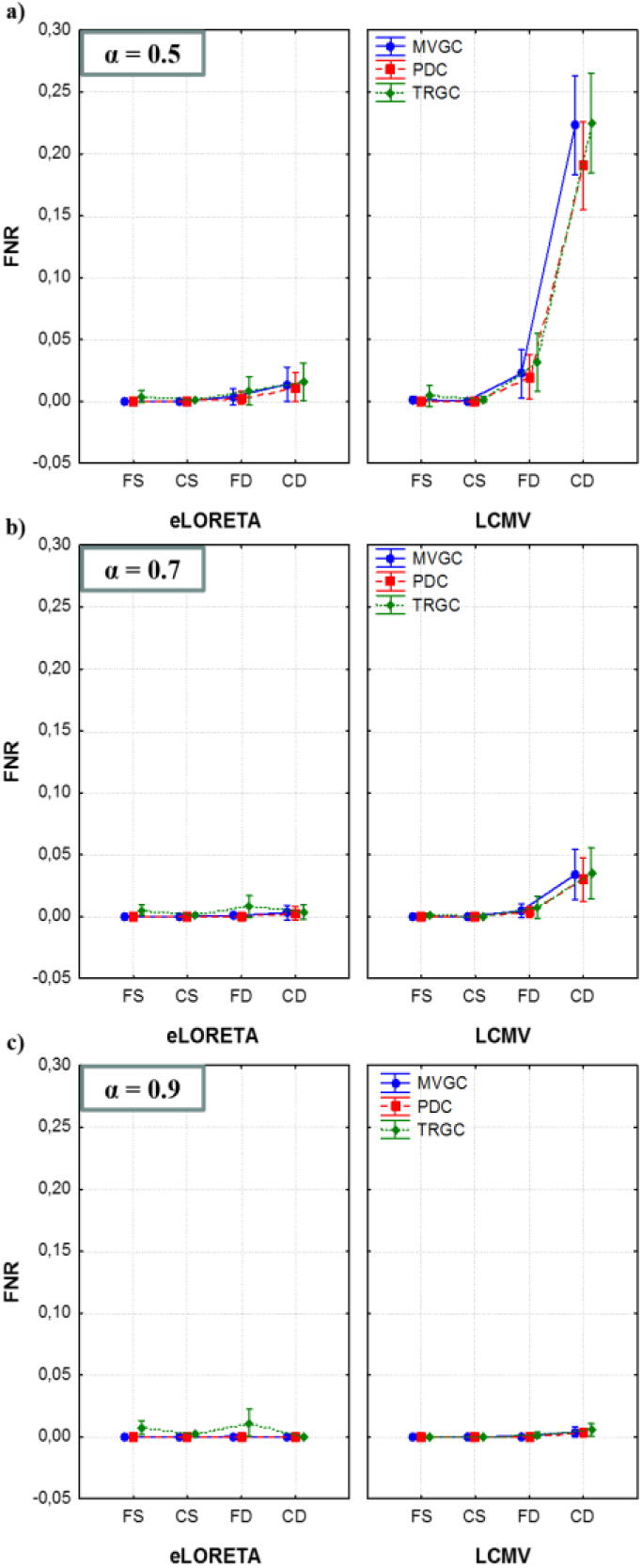
Means associated with the four-way interaction factor (L_INV METH x EST_TYPE x SNR x DIP_POS) of the ANOVA performed on the FNR. Each panel corresponds to a specific α level: 0.5 (panel a), 0.7 (panel b), 0.9 (panel c). For each panel, there are two graphs associated with the two different inverse solutions: eLORETA on the left and LCMV on the right. X-axes always show the levels of the factor DIP_POS and the colours code for the three connectivity estimators. Whiskers represent 95% confidence intervals. Each panel depicts results obtained for one SNR level.

The percentage of false negatives is less than 5% in all simulated cases, except for the lowest SNR level (α equal to 0.5) when LCMV is employed. In the easier condition with a higher signal to noise ratio and interacting dipoles that are not deep and close at the same time, the FNR is around 1% regardless of the chosen connectivity estimator.

##### Connectivity Estimator

The factor EST_TYPE does not have a significant effect on the FNR index independently of all the other factors (SNR value, type of algorithm chosen for the source reconstruction and connectivity estimation): its variations never exceed 1%. Also, the slight increase of false positives associated with the time reversed adaptation of GC is not statistically significant in this case.

##### Inverse Algorithm

The percentage of FN obtained with the two inverse methods is strictly linked to the dipoles’ position. Results reported in panel a) show that for SNR equal to 0db, FNR significantly increases for LCMV only when the dipoles are located deep in the brain (accounting for an increase of 20% in the *Close/Deep* condition). Panel b) shows a similar but attenuated trend for α equal to 0.7 (increase of less than 5% in the *Close/Deep* condition). As shown in panel c), there are no significant differences between LCMV and eLORETA for the highest SNR value.

##### Fixed dipoles position

The factor DIP_POS is significant for low and medium SNR values and L_INV METH corresponding to LCMV. In such conditions, for deep dipoles, the FNR is significantly higher, regardless the connectivity estimator. Moreover, focusing on the deep locations, there is a significant increase of the false negatives when the dipoles are close compared to when they are further away.

##### SNR

the signal-to-noise ratio associated to the three levels of the factor SNR significantly influences the presence of false negatives only when the inverse problem is solved by the LCMV algorithm. This is particularly the case for the condition *Close/Deep*, in which the FNR decreases from 20% when α is equal to 0.5 (panel a) to 4% when α is equal to 0.7 (panel b), and to 1% for the highest SNR level (panel c). This suggests that the amount of false negatives is independent of the algorithm employed for solving the inverse problem and for the connectivity estimation. In case of poor signal quality (low SNR), the FNR is considerable when the sources to be reconstructed are located deep in the brain. Algorithms that are more prone to missing connections are LCMV for source reconstruction and TRGC as connectivity estimation.

#### AUC

The graphs in Figure 5 depict the means of the four-way interaction factor (L_INV METH x EST_TYPE x SNR x DIP_POS) obtained for the AUC parameter.

**Figure 5.**
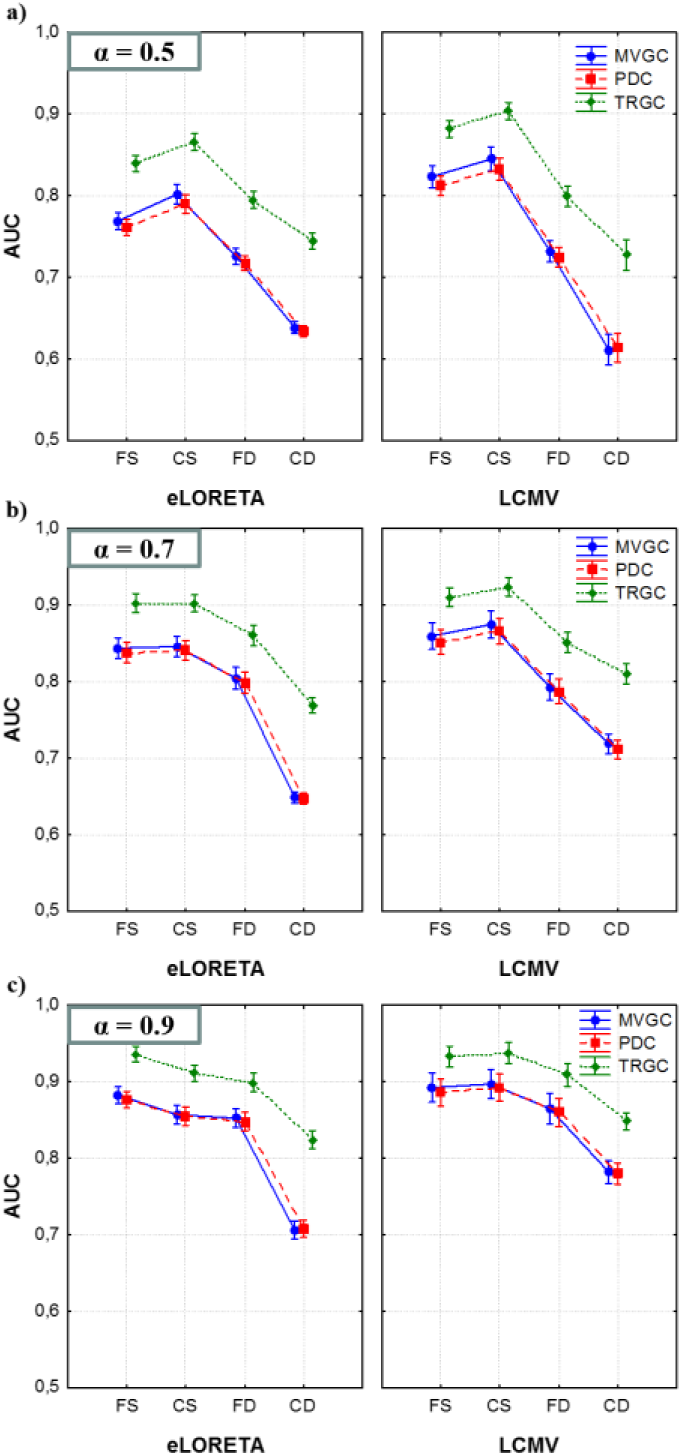
Means associated with the four-way interaction factor (L_INV METH x EST_TYPE x SNR x DIP_POS) of the ANOVA performed on the FNR. Each panel corresponds to a specific α level: 0.5 (panel a), 0.7 (panel b), 0.9 (panel c). In each panel, there are two graphs associated with the two different inverse solution: LCMV on the left and eLORETA on the right. X-axes report the levels of the factor DIP_POS and colours code for the three connectivity estimators. Whiskers represent 95% confidence intervals. Each panel depicts results obtained for one SNR level.

The AUC index hereby summarizes the effect of the four considered factors on the accuracy of the estimation in term of false positives and false negatives, providing a unifying measure of the discriminability of actually present and non-existent connections.

##### Connectivity Estimator

As expected from the previous results concerning the FPR trend, the accuracy of the estimation considerably increases when performed by means of TRGC. The increase of the performances associated with TRGC is statistically significant and amounts to about 10%. Confirming what has already been demonstrated for FPR and FNR, the accuracy of MVGC and PDC is not significantly different in any condition.

##### Inverse Algorithm

On average, the difference between LCMV and eLORETA is not significant, but there are combinations of the factors for which either of the two performed better. The main discrimination is given by the linked dipoles position. When the sources are located deep in the brain (especially if they are also close), the accuracy of the connectivity estimation appears significantly higher when LCMV is employed to reconstruct the brain activity. Once again, the only exception is the low SNR setting, in which this relationship is reversed because LCMV is more sensitive to the SNR level compared to the eLORETA algorithm, which shows more stable performance.

##### Fixed dipoles position

Independent of the employed inverse algorithm and connectivity estimator, the accuracy of the estimation significantly decreases when the linked dipoles are located deep in the brain. For higher SNR levels, the ANOVA highlights a performance degradation in terms of the AUC dropping from 90% (*Far/Superficial*) to 70% (*Close/Deep*).

##### SNR

As expected, the performance significantly improves in all considered conditions when the simulated SNR is high. More specifically, when the SNR level is 0.9, the accuracy is higher than 85% for eLORETA and higher than 90% for LCMV in all the cases except for the *Deep/Close* condition.

The analysis of the AUC index suggests that the optimal combination of factors is given by: i) dipoles located superficial in the brain and not too close; ii) LCMV algorithm when the SNR is not too low, otherwise eLORETA; iii) TRGC as connectivity estimator.

#### 3.2 Brain maps

##### Non-Interacting dipole position

As mentioned before, in each simulated condition, the moving dipole changes its position over 1004 locations equally distributed in the brain. In order to map the performance of the three connectivity estimators for each investigated source reconstruction algorithm and each position of the fixed dipoles, MVGC, PDC and TRGC were computed considering all the 1004 possible configurations of the network. Since each simulation was iterated 100 times, we were able to obtain an average performance value. Only the maps depicting the FPR are reported because of the greater sensitivity of this indicator to the factors considered in the analysis. We report transparent axial views of the head for each choice of fixed dipoles position and inverse method. Only 2 out of 3 estimators are reported because the PDC performances are similar to those obtained using the MVGC algorithm in all the considered conditions. We report the value assumed by the FPR (coded by its color) in the position of the moving dipole associated with that measure. Figure 6 reports the results obtained for the lowest SNR level, when α is equal to 0.5.

**Figure 6.**
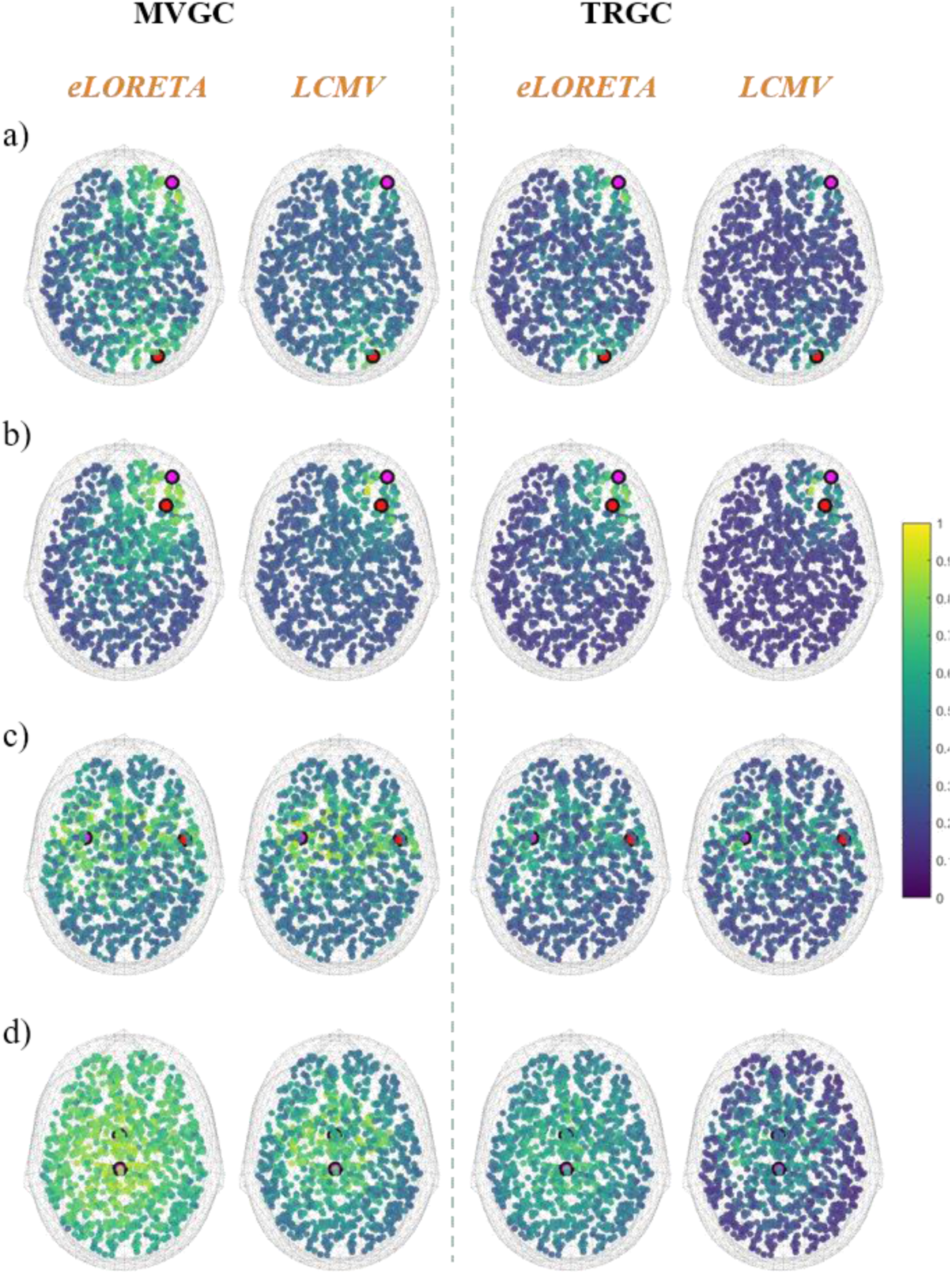
Spatial distribution of the FPR in the Moving Non-Interacting Dipole condition for low SNR (α = 0.5). Shown are the Sender (red circle) and Receiver (purple circle) of the interaction in the Far-Superficial (a), Far-Deep (b), Close-Superficial (c), and Close-Deep (d) conditions. The other points represent the mean value of the FPR across 100 iterations (coded by the colour bar on the right side) when the third active dipole (the Non-Interacting one) is moved across the brain. The first two columns refer to the classical GC (MVGC algorithm); the last two to the TRGC. For each column, results obtained with eLORETA and LCMV are reported next to each other.

The percentage of false positives depends on the distance of the Non-Interacting dipole from the two fixed ones. The most relevant result is that when the fixed dipoles are located deep in the brain and close to each other (panel d), high FPR values are spread across the whole brain, and reach 100% in the vicinity of the Sender and Receiver. Only TRGC combined with the LCMV algorithm mitigates this effect, which is then limited to the configurations in which the Non-Interacting dipole is close to the other two. Panels a), b) and c) clearly show a strong increase of the FPR when the Non-Interacting dipole is located in the areas close to the Receiver or to the Sender. Similar maps displaying the results obtained for α equal to 0.7 and 0.9 are reported in the supplementary material. These results confirm the trends commented for the previous maps but with globally better performances. The FPR considerably increases around the fixed dipoles. This phenomenon is focal when LCMV is employed and more spread-out if eLORETA combined with the MVGC estimator. Again, when the fixed dipoles are located deep in the brain and close each other, high FPR values are spread across the whole brain and reach 100% in the vicinity of Receiver and Sender. Maps associated with all the others fixed dipoles positions show that when the sources included in the model are far one from the other, the best performance is obtained with eLORETA. In order to summarize the information contained in these maps, figure 7 shows the value of the FPR as function of the distance of the moving dipole from the Sender of the interaction for all SNR values, inverse algorithms and connectivity estimators.

**Figure 7.**
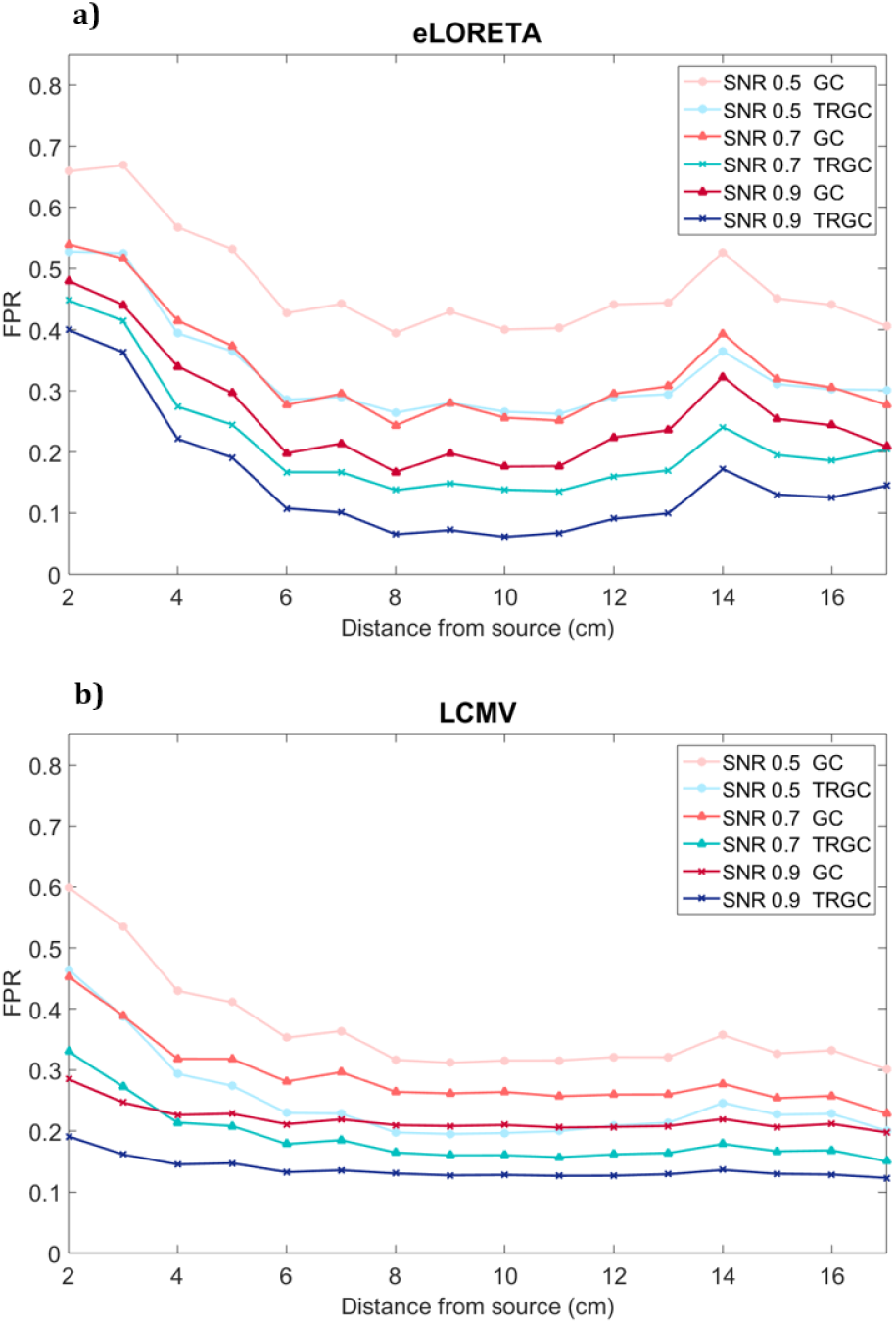
FPR as function of the distance (in cm) of the Non-Interacting Moving Dipole from the Sender of the interaction for the two inverse reconstruction algorithms, eLORETA and LCMV. The MVGC and TRGC connectivity estimators are drawn in red and blue colours, respectively. The circle marker codes for low SNR, the triangle for medium SNR, and the cross for high SNR.

The results suggest that TRGC performs better than MVGC regardless of the distance of the moving dipole from the Sender. LCMV source reconstruction is less sensitive to the distance between dipoles. For example, in panel a), an increase of the FPR 14 cm away from the Sender is noticeable. This point corresponds to the position of the second interacting dipole. When the LCMV algorithm is employed the increase is much less evident. The trends are similar for all α levels, although higher FPRs are observed for lower SNRs. For high SNR, the best performance is achieved with eLORETA when the moving dipole is far from the other two.

#### Interactive dipole position

The last analysis was performed using a fixed location for the Non-Interacting and Sender dipoles, placing the Receiver dipole at different positions. The results are in line with the previous ones. Figure 8 depicts topographical maps for the low SNR level (in the supplementary material for medium and high SNR levels), while figure 9 depicts FPR as a function of the distance between Receiver and Sender.

**Figure 8.**
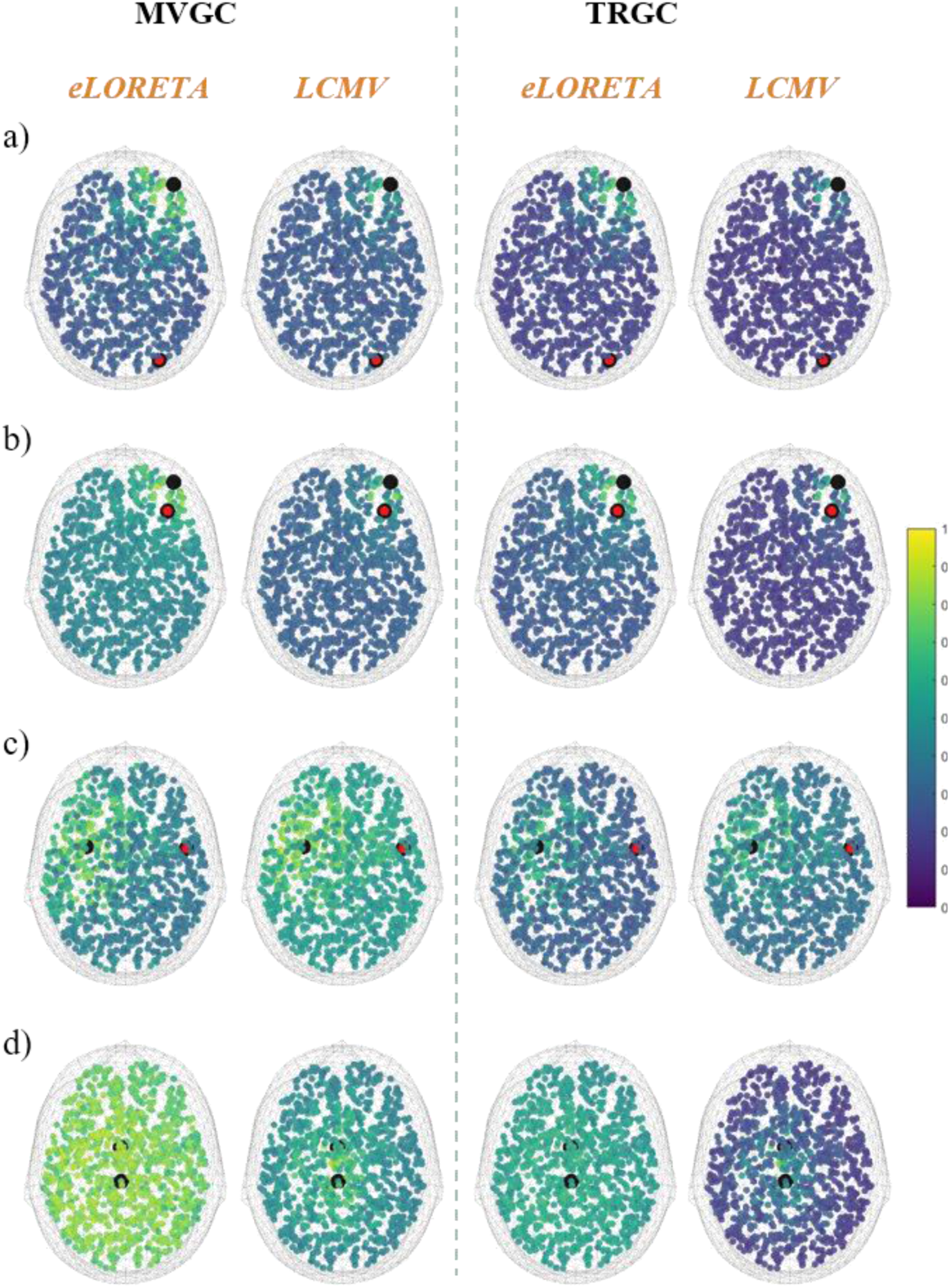
Spatial distribution of the FPR in the Moving Receiver Dipole condition for low SNR (α = 0.5). Shown are the Sender (red circle) and Receiver (purple circle) of the interaction in the Far-Superficial (a), Far-Deep (b), Close-Superficial (c), and Close-Deep (d) conditions. The other points represent the mean value of the FPR across 100 iterations (coded by the colour bar on the right side) when the third active dipole (the Non-Interacting one) is moved across the brain. The first two columns refer to the classical GC (MVGC algorithm); the last two to the TRGC. For each column, results obtained with eLORETA and LCMV are reported next to each other.

**Figure 9.**
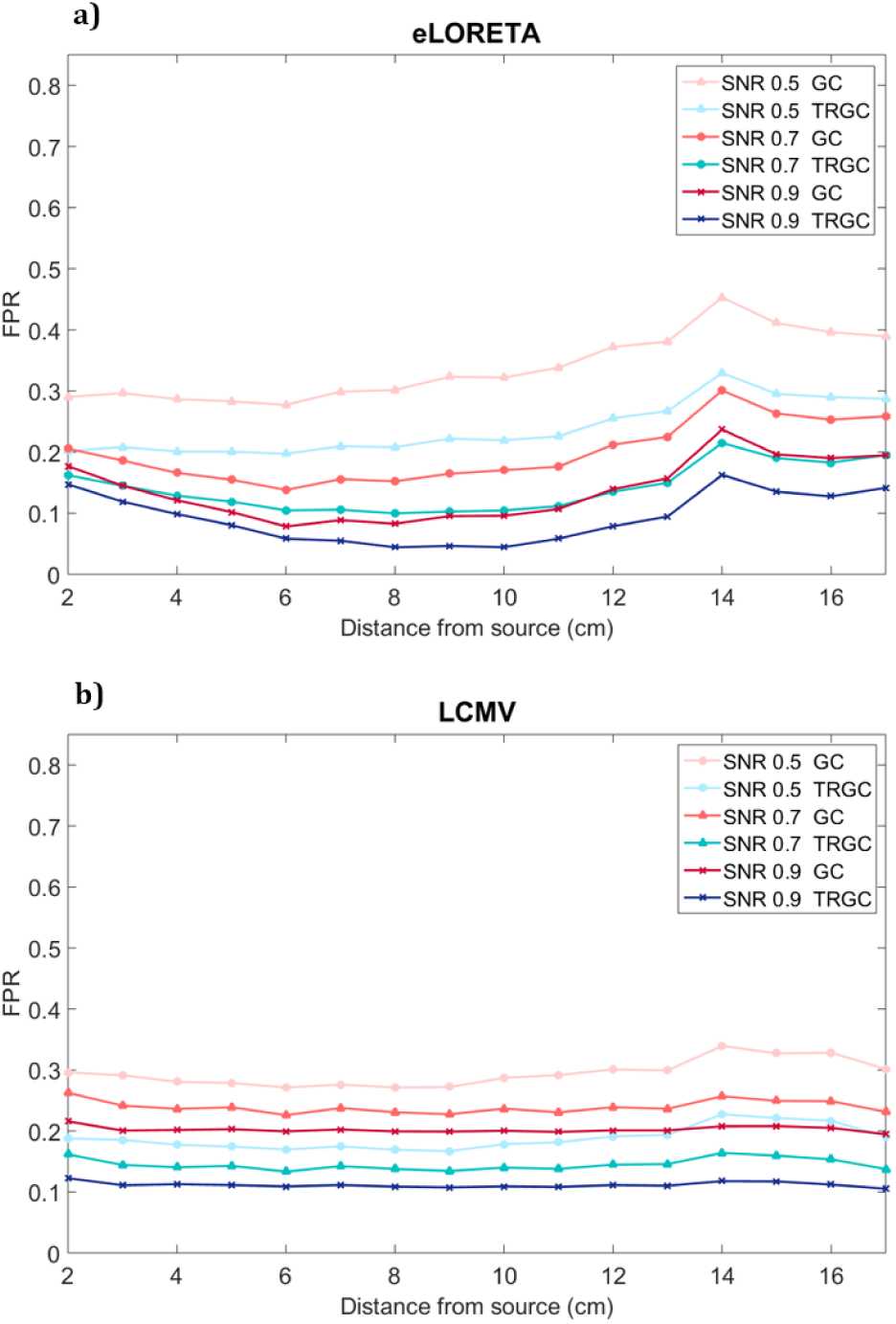
FPR as function of the distance (in cm) of the moving Receiver from the Sender for the two inverse reconstruction algorithms, eLORETA and LCMV. The MVGC and TRGC connectivity estimators are drawn in red and blue colours, respectively. The circle marker codes for low SNR, the triangle for medium SNR, and the cross for high SNR.

When the fixed dipoles are close (panels b and d), a high percentage of false positives appears throughout the brain, in particular for low and medium level of SNR. All others results are in line with the results reported above:

- the increase of the α value corresponds to a decrease in the number of false positives independently of all the other factors;
- on average, LCMV performed better than eLORETA. This advantage is predominantly due to an increased robustness w.r.t. the position of the nodes;
- TRGC provided more accurate connectivity estimates than MVGC and PDC;
- when the involved dipoles are far away from another, eLORETA leads to more accurate connectivity estimation, and the difference between the classical MVGC and TRGC is less pronounced than in the other conditions;
- with the dipoles in the *Far/Superficial* configuration, and α equal to 0.9, the percentage of false positives is less than 10% for all the inverse solutions and connectivity algorithms;
- with the dipoles in the *Close/Deep* configuration, the percentage of false positives reaches 100% regardless of the SNR value.

Fig. 9 shows the FPR as function of the distance between Sender and Receiver for all SNR values, inverse algorithms and connectivity estimators.

The first result is that the mean value of the FPR is lower than for the Non-Interacting moving dipole condition. Also, in this case, TRGC performed better than MVGC regardless of the distance of the moving Receiver dipole from the Sender. In panel a) it is possible to notice an increase of the FPR when the Receiver dipole is 14cm away from the Sender dipole (this being the position of the Non-Interacting dipole). Trends are similar for all the α levels, where, generally, decreases in SNR are associated with increases in FPR. For high SNR, the best performance is achieved using eLORETA when the moving dipole is far away from the other two (FPR around 5%).

## 4. Discussion and Conclusion

It is well established that neuroelectrical measures recorded on the scalp need to be projected back into the brain in order to be able to infer at least roughly where these signals have been generated. In the same way it is evident that measures of statistical dependencies between brain regions cannot be inferred by studying dependencies between scalp sensor signals [8]–[10]. Unfortunately, even with state-of-the-art localization of the brain sources underlying the measured signals, directed dynamical influences between these reconstructed sources do not always reflect the ground truth. This issue has been anticipated in [8]–[10] and thoroughly analyzed by Palva and colleagues [43] for phase-based (undirected) connectivity measures. In the present comprehensive simulation study, we focused on directional connectivity measures and quantified the extent to which the estimation of influences between reconstructed sources is possible. We employed an analysis framework combining source localization approaches and brain connectivity estimators with the goal of identifying those analysis pipelines that that are least affected by the presence of head volume conduction and, therefore, provide the most accurate and reliable connectivity estimates. Several realistic conditions of brain activity were simulated, where our goal was to simulate both advantageous and disadvantageous conditions for the following brain connectivity estimation. To this end, we modulated the depth of the sources, the distance between sources, and the SNR. Not surprisingly, a convenient condition we identified is the presence of far and superficial dipoles in combination with a high SNR; in contrast, a disadvantageous condition is given by the presence of close and deep sources with a low SNR level. Our simulations suggest that all considered factors show a significant influence on the estimation quality and, consequently, their combination has a considerable impact on the connectivity estimation performance. LCMV source reconstruction appears to be more sensitive to the SNR value, while eLORETA achieves similar performance regardless of the SNR. In general, LCMV showed better performance than eLORETA. Only when the simulated sources were assigned to distant locations, the eLORETA performance is similar to or better than the performance of LCMV. In agreement with the theoretical hypothesis, we demonstrated that the TRGC algorithm provides a better estimation of the directed statistical dependencies between sources than classical MVGC and PDC. Indeed, the percentage of spurious connections decreased significantly and the overall detection of connectivity as measured by the AUC increased significantly in all considered experimental conditions when TRGC was used instead of MVGC or PDC. At the same time, the percentage of missed connections as measured by the FNR increased slightly, but still remained close to zero. GC and PDC showed similar performance independent of all other factors. As expected, we found that closer and deeper active sources decreased the obtained performance. Thus, a dependence between the dipoles position and the accuracy of the estimates was found. This is a clear effect of the volume conduction, since, when two sources are close to each other, they generate a highly mixed signal on the scalp, which compromises the correct estimation even after inverse source reconstruction. On the other hand, when the sources are far away from each other, they are less affected by volume conduction, leading to a better quality of the connectivity estimation. The insights obtained in this study may guide the choice of crucial parameters such as selection of regions-of-interest (ROI) as well as the selection of source reconstruction and connectivity estimation algorithms that promise to provide the most reliable and physiologically interpretable description of brain networks based on EEG data.

We agree with [43], advocating for the application of measures, for which promises and pitfalls are known, and which integrate knowledge of how neural activity in the whole brain as well as external (physiological or artifactual) activity contribute to the signals that we record on the scalp. In this regard, it should be noted that connectivity estimates can only at most be as focal as the reconstructed source current densities they are derived from, and we know that common inverse methods lead to very blurry results. To distinguish correctly-identified connections from connections that are observed in the vicinity of the true interacting sources due to blurry inverse solutions, a data-driven clustering in the space of brain-wide pairwise connectivities, as recently proposed in [44], may a viable option, which may be preferable to a reduction of the source space to the level of static ROIs. It has to be kept in mind, however, that – although of importance – the main problem in EEG-based brain connectivity analysis is not the spatial blur of correctly identified connections but the emergence of spurious connectivity as a result of observing mixtures of signals even at the level of reconstructed sources. This problem can only be addressed by using appropriate connectivity measures that are robust to volume conduction effects by construction.

### Code and data availability

The code necessary to reproduce these simulations is available at: https://github.com/paolop21/simulation_source_connectivity.

The results of the simulations and the structures necessary to run the code are available at: https://zenodo.org/record/1155857#.WmMVwqjiY2w

